# Absence of biofilm adhesin proteins changes surface attachment and cell strategy for *Desulfovibrio vulgaris* Hildenborough

**DOI:** 10.1101/2024.09.10.612303

**Authors:** C. Pete Pickens, Dongyu Wang, Chongle Pan, Kara B. De León

**Author notes:** Corresponding Author: Kara B. De León, 770 Van Vleet Oval Norman, Oklahoma, USA.

## Abstract

Ubiquitous in nature, biofilms provide stability in a fluctuating environment and provide protection from stressors. Biofilms formed in industrial processes are exceedingly problematic and costly. While biofilms of sulfate-reducing bacteria in the environment are often beneficial because of their capacity to remove toxic metals from water, in industrial pipelines, these biofilms cause a major economic impact due to their involvement in metal and concrete corrosion. The mechanisms by which biofilms of sulfate-reducing bacteria form, however, is not well understood. Our previous work identified two proteins, named by their gene loci DVU1012 and DVU1545, as adhesins in the model sulfate-reducing bacterium, *Desulfovibrio vulgaris* Hildenborough. Both proteins are localized to the cell surface and the presence of at least one of the proteins, with either being sufficient, is necessary for biofilm formation to occur. In this study, differences in cell attachment and early biofilm formation in single deletion mutants of these adhesins were identified. Cells lacking DVU1012 had a different attachment strategy from wild-type and ΔDVU1545 cells, more often attaching as single cells than aggregates, which indicated that DVU1012 was more important for cell-to-cell attachment. ΔDVU1545 cells had increased cell attachment compared to wild-type cells when grown in static cultures. To date, comparisons of the *D. vulgaris* Hildenborough have been made to the large adhesion protein (Lap) system in environmental pseudomonads. Yet, we and others have shown distinct mechanistic differences in the systems. We propose to name these proteins in *D. vulgaris* Hildenborough biofilm formation system (Bfs) to facilitate comparisons.

**Importance:** Biofilms of sulfate-reducing bacteria contribute to biocorrosion, costing the U.S. hundreds of millions of dollars annually. In contrast, these biofilms can be used to bioremediate toxic heavy metals and to generate bioelectricity. As one of the most abundant groups of organisms on Earth, it is pertinent to better understand mechanistically how the biofilms of sulfate-reducing bacteria form so we may use this knowledge to help in efforts to mitigate biocorrosion, to promote bioremediation, and to produce clean energy. This study shows that the absence of either one of two biofilm adhesins impacts surface colonization by a sulfate-reducing bacterium, and that these two biofilm adhesins differ in their effect on cell attachment compared to other well-documented bacteria such as *Pseudomonas* species.

## Introduction

Ubiquitous in nature, biofilms are densely packed microbial cells attached to surfaces that envelop themselves in a matrix of extracellular polymeric substance (EPS) made of carbohydrates, proteins, DNA, and other polymers. Growth as a biofilm is evolutionarily advantageous as it gives some stability in a constantly fluctuating environment and provides protection from a variety of environmental challenges including UV, acid, dehydration, salinity, metal toxicity, phagocytosis, and several antimicrobial agents (1). In subsurface environments, the active microbial population is predominantly found attached to surfaces as a biofilm (2).

Biofilms of sulfate-reducing bacteria (SRB) impact the world around them in multiple ways. SRB biofilms are the major contributors to microbiologically influenced corrosion (MIC), causing millions of dollars in damages annually to the United States (3). SRB biofilms can also assist with bioremediation, where SRB biofilms precipitate heavy metals such as chromium and uranium and allow for their removal from contaminated groundwater (4, 5). These environmental effects caused by SRB biofilm showcase their impact and importance in industrial and environmental systems. Biofilms of SRB have also been shown to play a role in animal gut colonization and have been shown to affect immune responses in hosts through the production of sulfide gas (6). These effects and outcomes of SRB biofilm have been well-documented, affecting multiple different environments and having potentially positive and negative effects in their immediate environment.

Biofilms formed by the model SRB, *Desulfovibrio vulgaris* Hildenborough (7), have previously been shown to be predominantly protein and are a distinct state of growth for *D.* vulgaris cells compared to cells growing unassociated with a surface (8, 9). Previous work identified the importance of a type I secretion system and two biofilm adhesins in biofilm formation under continuous culture and flow (10). Those proteins are annotated as hemolysin-like calcium binding proteins and are called by their gene loci, DVU1012 and DVU1545. The absence of either one of these two proteins caused no difference in biofilm formation over time compared to the wild-type strain. However, in the absence of both proteins no biofilm was formed.

There are other examples of biofilm adhesins affecting surface colonization and biofilm maturation across other bacterial species. In the bacteria *Pseudomonas fluorescens* and *P.* putida, proteins LapA and LapF share similarities to *D. vulgaris* proteins DVU1012 and DVU1545 (11, 12). Both proteins have been well-characterized for their role in surface colonization and biotic attachment. LapA and LapF also utilize a type I secretion system (LapBC and TolC) for export out of the cell, and both are retained in the outer membrane. However, mutants lacking the protein LapA was found to be deficient in surface colonization and whole biofilm formation over time compared to LapF, whereas the protein LapF was found to have a greater impact on cell-to-cell adhesion and later stage biofilm development (11, 12). Retention of the protein LapA on the surface of the cell is determined by the activity of the periplasmic protease LapG, which in turn is regulated by the cyclic-di-GMP sensing protein LapD; no such retention regulation mechanism has been shown for the protein LapF. DVU1019 and DVU1020 in *D. vulgaris* Hildenborough have been shown to perform similar functions to LapG and LapD, respectively (13). These proteins have served as examples for biofilm formation for homolog adhesins across other species and have shown how multiple biofilm adhesins within a host cell can perform different functions.

For *D. vulgaris* Hildenborough biofilm proteins DVU1012 and DVU1545, their distinct functional roles have not yet been determined. The protein DVU1012 was previously found to be the most abundant protein within the extracellular matrices of the biofilm and has been shown to have high abundance within planktonic cultures as well (9, 14). The other biofilm-associated protein DVU1545, however, is found in lower abundance compared to DVU1012. Since the absence of either DVU1012 or DVU1545 does not impact biofilm formation, biofilms formed by *D. vulgaris* must then compensate for the absence of one of the most abundant proteins within the extracellular biofilm matrix. As well, the absence of DVU1012 does not influence biofilm maturation over time, unlike the mutation of the analogous protein LapA in *P. fluorescens* and *P. putida* (11). As the proteins LapA and LapF have been shown to have distinct roles for biofilm growth in environmental *Pseudomonas* species, the proteins DVU1012 and DVU1545 may also play differing roles in surface colonization and maturation. While the consequences and outcomes of SRB biofilms are well understood, the mechanisms used by SRB to colonize surfaces, co-aggregate, and form biofilm are not. As complete redundancy of proteins is rare in microbes (15), we hypothesize that the two proteins DVU1012 and DVU1545 have distinct, but overlapping roles in cell attachment and biofilm formation. In this study, proteomics was used to assess whether the absence of either biofilm adhesin caused a proportional change in the other adhesin or in any other proteins within the biofilm matrix. Early static biofilms were quantified to determine differences in biofilm maturation at different development stages and under differing environmental conditions. Surface colonization and cell aggregation of *D. vulgaris* adhesin protein mutants were assessed using real-time phase contrast microscopy. This study highlights the effects of the biofilm adhesin deletion on mature biofilms of *D. vulgaris* Hildenborough and the major differences between the functional roles of SRB adhesins DVU1012 and DVU1545 compared to LapA and LapF in the environmental pseudomonads.

## Results

### Abundance of adhesin proteins DVU1012 or DVU1545 does not change in mutant biofilms lacking the other adhesin

Monitoring biofilm growth over time, no significant differences in total biomass were measured between the single deletion strains of DVU1012, DVU1545, and the parental strain, with minimal differences in measured proteins or hexose per square centimeter (Figure 1). This corresponds with previous findings with protein measurements from *D. vulgaris* Hildenborough biofilms (10). Carbohydrate measurements for biofilms of the single deletion mutant strains have not previously been published, and no significant differences were found between the strains. Since there was no difference in measured biofilm protein between the strains and the biofilms were still predominantly protein, proteomic analyses were performed to determine differences in biofilm protein composition. Whole biofilm samples from 72 h for each strain were used in proteomic analyses (Table S1). Comparing normalized protein intensities, which are on a log_2_-based scale, DVU1012 was approximately four-fold more abundant than DVU1545 in wild-type cells (Figure 2a). Overall, the absence of either biofilm adhesin does change the biofilm protein compositions between the single deletion mutants and wild-type (WT) cultures (Figure 2b-c). Interestingly, when comparing the relative protein abundance of the proteins DVU1012 and DVU1545 between WT and the single deletion mutants, the abundance of the proteins DVU1012 and DVU1545 was not different between WT and the corresponding single deletion mutants (Figure 2a). In addition, few other proteins changed in abundance between WT and the single deletion mutants (Figure 2b-c). Biofilms formed by the ΔDVU1012 strain had decreased abundance of proteins related to cell wall/membrane/envelope biogenesis and coenzyme transport and metabolism (Table 1). Most proteins with different abundance in biofilms of the ΔDVU1012 strain when compared to WT were predicted to be localized to the cytoplasm, and none of the proteins were predicted to be localized to the inner or outer membrane. This differs when looking at the proteins with different abundances in ΔDVU1545 biofilms compared to WT, as four proteins are predicted to localize to the inner membrane (Table 2). Proteins related to cell wall/membrane/envelope biogenesis were increased overall in ΔDVU1545 biofilms compared to WT, and proteins predicted to be involved in coenzyme transport and metabolism were decreased. None of the proteins with different abundances in the single deletion strains compared to WT had annotations that supported a role in biofilm matrix formation. Thus, overall, the absence of either biofilm adhesin protein did not cause a significant change in abundance of the other protein, and another protein did not apparently compensate for the missing adhesin.

**Figure 1.**
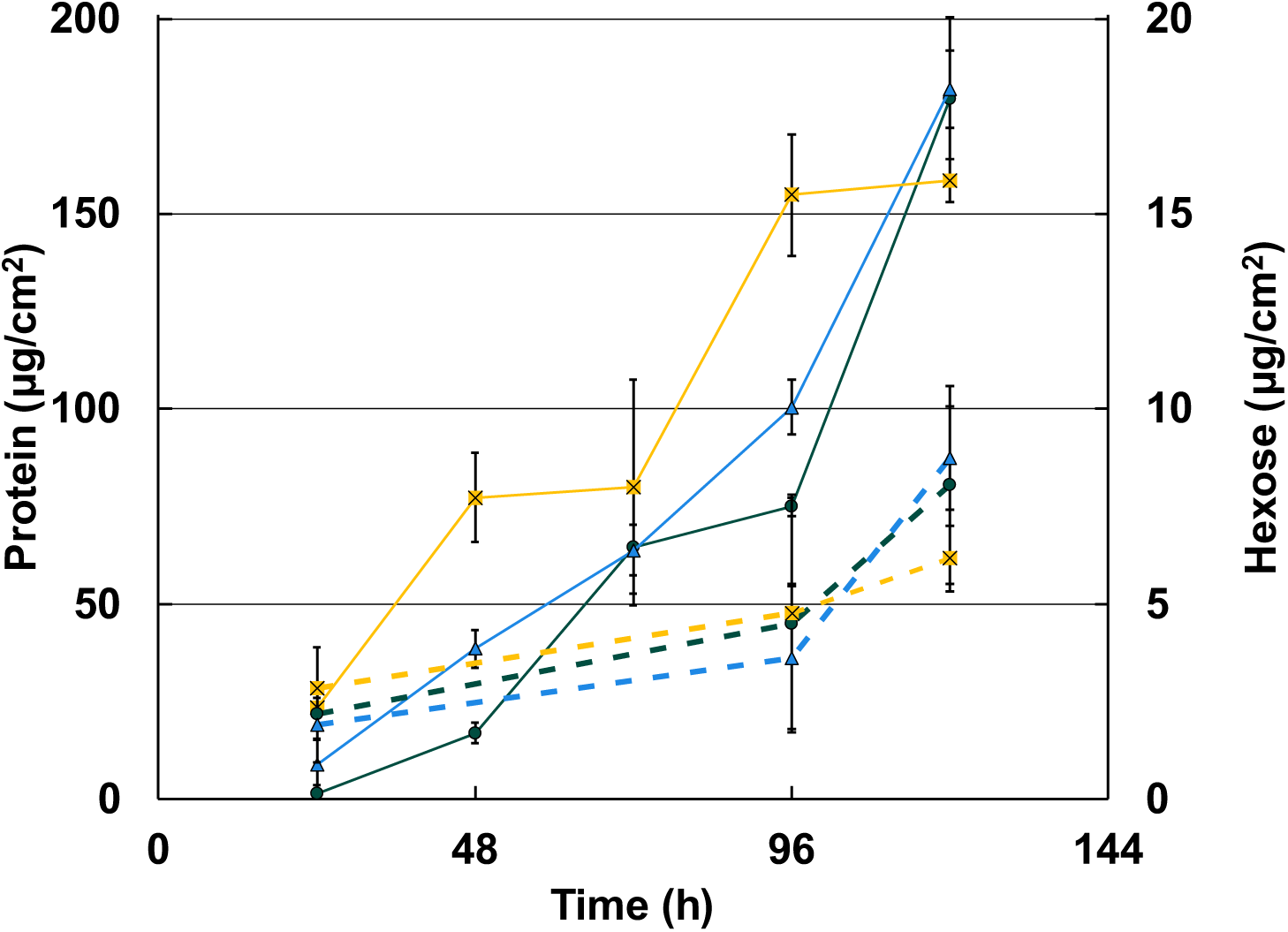
Biofilm maturation over time for WT and the biofilm adhesin mutants. Protein (solid lines) and hexose (dashed lines) as micrograms per square centimeter over time for WT (dark teal, circle), ΔDVU1545 (yellow, square), and ΔDVU1012 (blue, triangle). Error bars indicate standard deviation across technical replicates (n=3).

**Figure 2.**
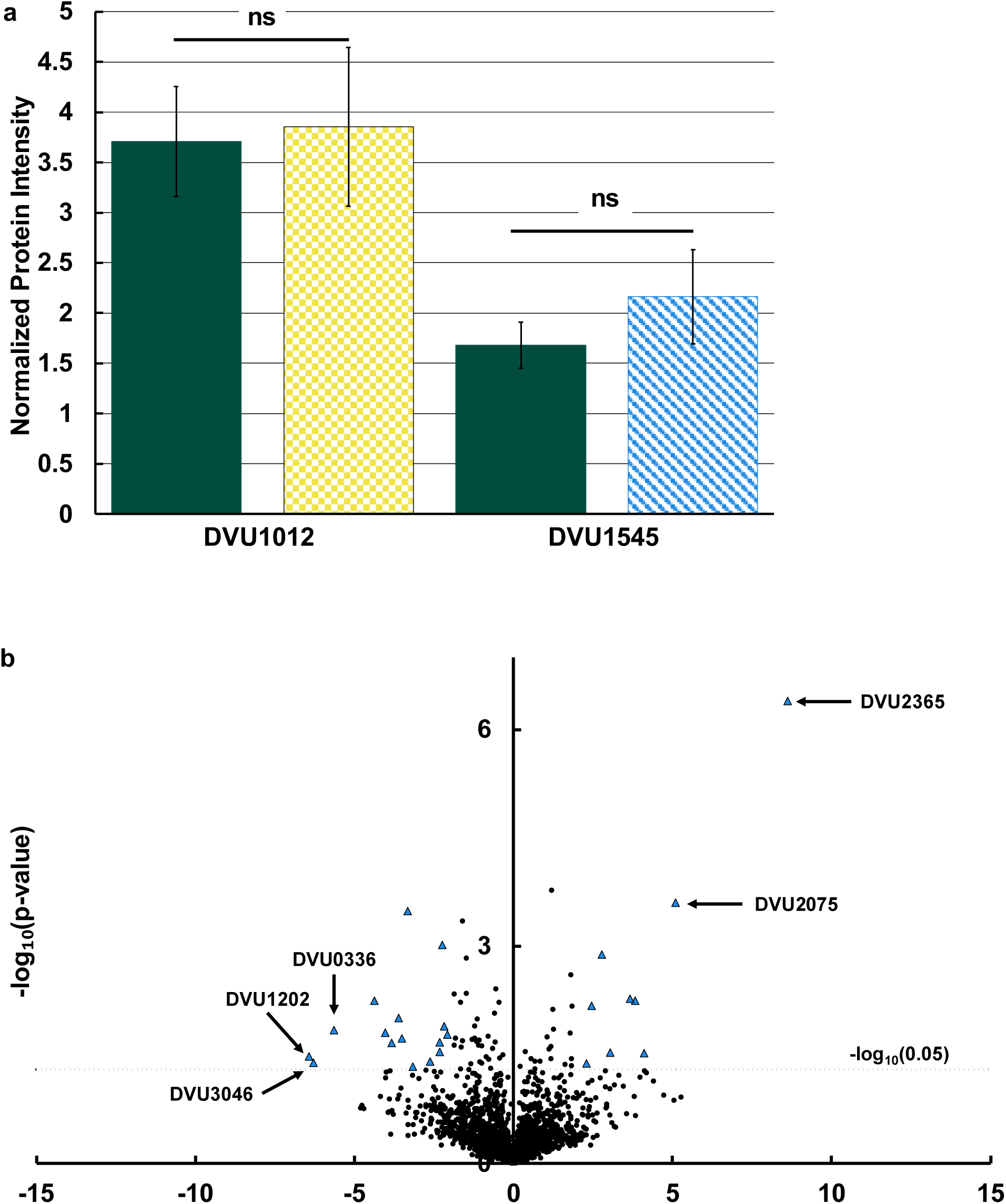
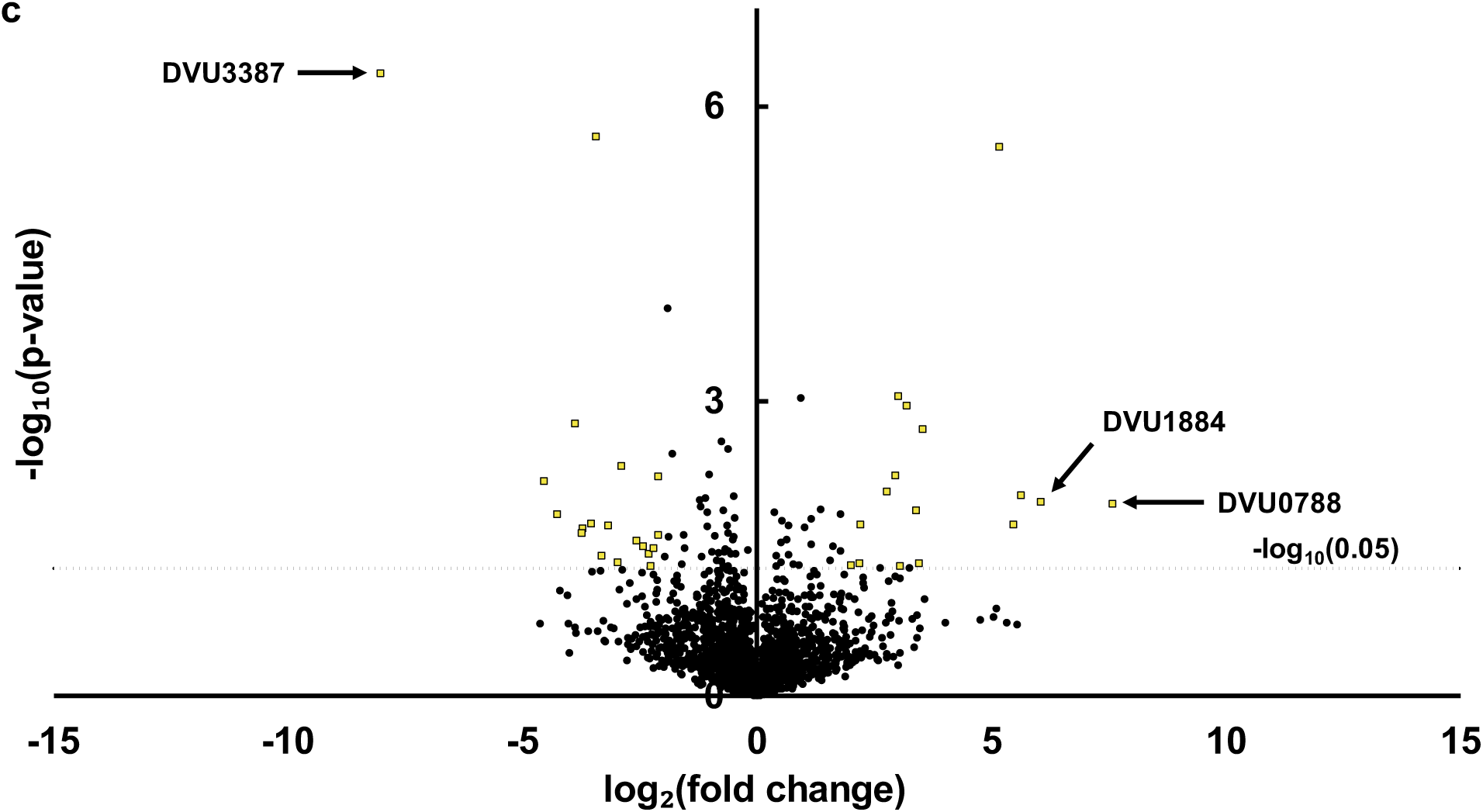
Changes in protein abundance between biofilms of WT and the biofilm adhesin mutants. (a) Normalized protein intensity for DVU1012 and DVU1545 for WT (dark teal, solid); ΔDVU1545 (yellow, checker); and ΔDVU1012 (blue, slant). Error bars indicate standard deviation across replicates (n=3). ns; non-significant differences as determined by a homoscedastic Student’s t-test (p > 0.05). The relative fold change in abundance of proteins were compared between WT and ΔDVU1012 (b) and WT and ΔDVU1545 (c). Dotted line denotes the cut-off for statistical significance (p < 0.05). Data points with larger than a four-fold change in abundance and statistically significant are marked for ΔDVU1012 (blue, triangle) and ΔDVU1545 (yellow, square). The gene locus of selected proteins are labeled with an arrow. The complete lists of significant proteins are shown in Tables 1 and 2.

**Table 1.** Proteins with significant fold change differences between ΔDVU1012 and WT.

| Gene locus ID | Description | Functional code<sup>a</sup> | Predicted Localization<sup>b</sup> | Fold Change | p-value |
| --- | --- | --- | --- | --- | --- |
| DVU2365 | conserved hypothetical protein | S | Cytoplasm | 8.615 | 4.01E-07 |
| DVU2075 | ParA family protein | D | Cytoplasm | 5.104 | 2.47E-04 |
| DVU2135 | hypothetical protein |  | Extracellular | 4.114 | 0.030 |
| DVU3165 | conserved domain protein |  | Cytoplasm | 3.822 | 0.006 |
| DVU0573 | creA protein | S | Unknown | 3.667 | 0.005 |
| DVU2057 | conserved hypothetical protein | S | Cytoplasm | 3.031 | 0.029 |
| DVU0959 | replicative DNA helicase | L | Cytoplasm | 2.787 | 0.001 |
| DVU2577 | DNA-binding response regulator, LuxR family | TK | Cytoplasm | 2.463 | 0.007 |
| DVU1980 | hypothetical protein |  | Unknown | 2.288 | 0.042 |
| DVU1200 | riboflavin synthase, alpha subunit | H | Cytoplasm | -2.088 | 0.017 |
| DVU0984 | tRNA-i(6)A37 modification enzyme MiaB | J | Cytoplasm | -2.169 | 0.013 |
| DVUA0019 | type II DNA modification methyltransferase, putative |  | Cytoplasm | -2.239 | 0.001 |
| DVU0829 | phosphoenolpyruvate-protein phosphotransferase | G | Cytoplasm | -2.316 | 0.021 |
| DVU0469 | N-(5-phosphoribosyl)anthranilate isomerase | E | Cytoplasm | -2.325 | 0.028 |
| DVU1420 | Hpt domain protein | T | Unknown | -2.619 | 0.039 |
| DVU3227 | flagellar basal body-associated protein, putative | N | Unknown | -3.163 | 0.046 |
| DVU1596 | protein-glutamate methylesterase CheB | NT | Cytoplasm | -3.326 | 3.22E-04 |
| DVU0282 | A/G-specific adenine glycosylase | L | Cytoplasm | -3.501 | 0.019 |
| DVU1365 | heme-binding protein, putative | H | Unknown | -3.612 | 0.010 |
| DVU2748 | precorrin-4 C11-methyltransferase | H | Cytoplasm | -3.823 | 0.021 |
| DVU0135 | conserved hypothetical protein | D | Cytoplasm | -4.027 | 0.016 |
| DVU3265 | tartrate dehydratase beta subunit, putative | C | Cytoplasm | -4.379 | 0.006 |
| DVU0336 | aminotransferase, DegT/DnrJ/EryC1/StrS family | M | Cytoplasm | -5.638 | 0.014 |
| DVU3046 | glycosyl transferase, group 1 family protein | M | Cytoplasm | -6.283 | 0.040 |
| DVU1202 | cytidine/deoxycytidylate deaminase family protein | F | Unknown | -6.439 | 0.033 |

<sup>a</sup> Functional codes based on Clusters of Orthologous Groups (COGs).
<sup>b</sup> Localization of proteins predicted by PSORTb.

**Table 2.** Proteins with significant fold change differences between ΔDVU1545 and WT.

| Gene locus ID | Description | Functional code<sup>a</sup> | Predicted Localization<sup>b</sup> | Fold-change | p-value |
| --- | --- | --- | --- | --- | --- |
| DVU0788 | rod shape-determining protein MreC | M | Unknown | 7.584 | 0.011 |
| DVU1884 | methyl-accepting chemotaxis protein | NT | Inner Membrane | 6.066 | 0.011 |
| DVU1165 | NADH respiratory dehydrogenase | C | Inner Membrane | 5.639 | 0.009 |
| DVU2057 | conserved hypothetical protein | S | Cytoplasm | 5.481 | 0.018 |
| DVU2859 | tail sheath protein, putative | R | Unknown | 5.177 | 2.59E-06 |
| DVU3165 | conserved domain protein |  | Cytoplasm | 3.529 | 0.002 |
| DVU0888 | response regulator | T | Cytoplasm | 3.449 | 0.045 |
| DVU0867 | aromatic amino acid decarboxylase, putative | E | Cytoplasm | 3.406 | 0.013 |
| DVU0959 | replicative DNA helicase | L | Cytoplasm | 3.187 | 0.001 |
| DVU1889 | phosphoheptose isomerase | G | Cytoplasm | 3.048 | 0.048 |
| DVU0573 | creA protein | S | Unknown | 3.022 | 0.001 |
| DVU1651 | conserved hypothetical protein |  | Inner Membrane | 2.964 | 0.006 |
| DVU2063 | conserved hypothetical protein | L | Cytoplasm | 2.773 | 0.008 |
| DORF41366 | Hypothetical protein |  | Unknown | 2.204 | 0.018 |
| DVU1360 | UDP-glucose 4-epimerase | M | Cytoplasm | 2.186 | 0.045 |
| DVU2512 | S-adenosyl-methyltransferase MraW | M | Cytoplasm | 2.013 | 0.047 |
| DVU2070 | TPR domain protein | S | Unknown | -2.102 | 0.023 |
| DVU3223 | aspartate aminotransferase | E | Cytoplasm | -2.107 | 0.006 |
| DVU0091 | conserved hypothetical protein |  | Unknown | -2.211 | 0.032 |
| DVU0606 | transcriptional regulator, ArsR family/methyltransferase, UbiE/COQ5 family | H | Cytoplasm | -2.256 | 0.048 |
| DVU1346 | exodeoxyribonuclease VII, large subunit | L | Cytoplasm | -2.300 | 0.036 |
| DVU0401 | hypothetical protein |  | Unknown | -2.421 | 0.030 |
| DVU0659 | hypothetical protein |  | Unknown | -2.567 | 0.027 |
| DVU3310 | ATP-dependent RNA helicase, DEAD/DEAH family | LKJ | Cytoplasm | -2.895 | 0.005 |
| DVU1423 | 2-oxoglutarate dehydrogenase, E3 component, lipoamide dehydrogenase | C | Cytoplasm | -2.970 | 0.044 |
| DVU1775 | 3,4-dihydroxy-2-butanone 4-phosphate synthase | H | Cytoplasm | -3.165 | 0.018 |
| DVU0336 | aminotransferase, DegT/DnrJ/EryC1/StrS family | M | Cytoplasm | -3.318 | 0.037 |
| DVU0821 | conserved hypothetical protein |  | Unknown | -3.428 | 2.04E-06 |
| DVU2062 | ATP-dependent DNA helicase, UvrD/REP family | L | Cytoplasm | -3.529 | 0.018 |
| DVU1559 | aldehyde oxidoreductase | C | Cytoplasm | -3.708 | 0.020 |
| DVU2655 | D-alanyl-D-alanine carboxypeptidase family protein | M | Inner Membrane | -3.733 | 0.022 |
| DVU3049 | hemerythrin family protein | P | Cytoplasm | -3.874 | 0.002 |
| DVU1365 | heme-binding protein, putative | H | Unknown | -4.265 | 0.014 |
| DVU3387 | hypothetical protein |  | Unknown | -4.544 | 0.007 |

<sup>a</sup> Functional codes based on Clusters of Orthologous Groups (COGs).
<sup>b</sup> Localization of proteins predicted by PSORTb.

### Absence of protein DVU1545 increases biofilm formation under batch culture and static conditions

Our previous experiments comparing whole biofilm growth over time of WT and single deletion mutants in CDC bioreactors did not begin measurements of biofilm biomass until 48 h post-inoculation, following 24 h of batch growth and 24 h of continuous flow (10). At this timepoint, the ΔDVU1545 strain consistently had more biofilm biomass than WT and the biofilm of ΔDVU1012 strain was often not detectable. Yet, all strains formed similar amounts of biofilm during maturation. To determine whether the absence of either one or both biofilm adhesins affected early biofilm growth; biofilm samples from wild-type, single deletion mutants, and the double deletion mutant cultured under batch, static conditions within Balch tubes were compared after 24 h of growth (Figure 3). There were no significant differences in growth rate between the strains (growth rate constants; h^-1^ +/- standard deviation: WT, 0.16 +/- 0.01; ΔDVU1545, 0.17 +/- 0.01; ΔDVU1012, 0.17 +/- 0.01; and ΔDVU1012 ΔDVU1545, 0.16 +/- 0.01), and so differences in biofilm biomass would not be from differences in growth rate but due to the effect of the deletion of DVU1012 and/or DVU1545. Biofilms were grown statically in batch cultures for 24 h (approximately six doublings) as this allowed for biofilm biomass to be quantifiable, as 16 h was the earliest that biofilm biomass could reliably be measured by protein in the WT cultures. At 24 h, the cells would be in log phase and, thus, not considered to be nutrient limited under these culturing conditions. The ΔDVU1545 strain had increased biofilm compared to all other strains (Figure 3). The ΔDVU1012 strain had less biofilm compared to both WT and ΔDVU1545. The double deletion mutant had little to no biofilm formation, consistent with our previous findings from culturing biofilms under continuous culture and shear conditions (10).

**Figure 3.**
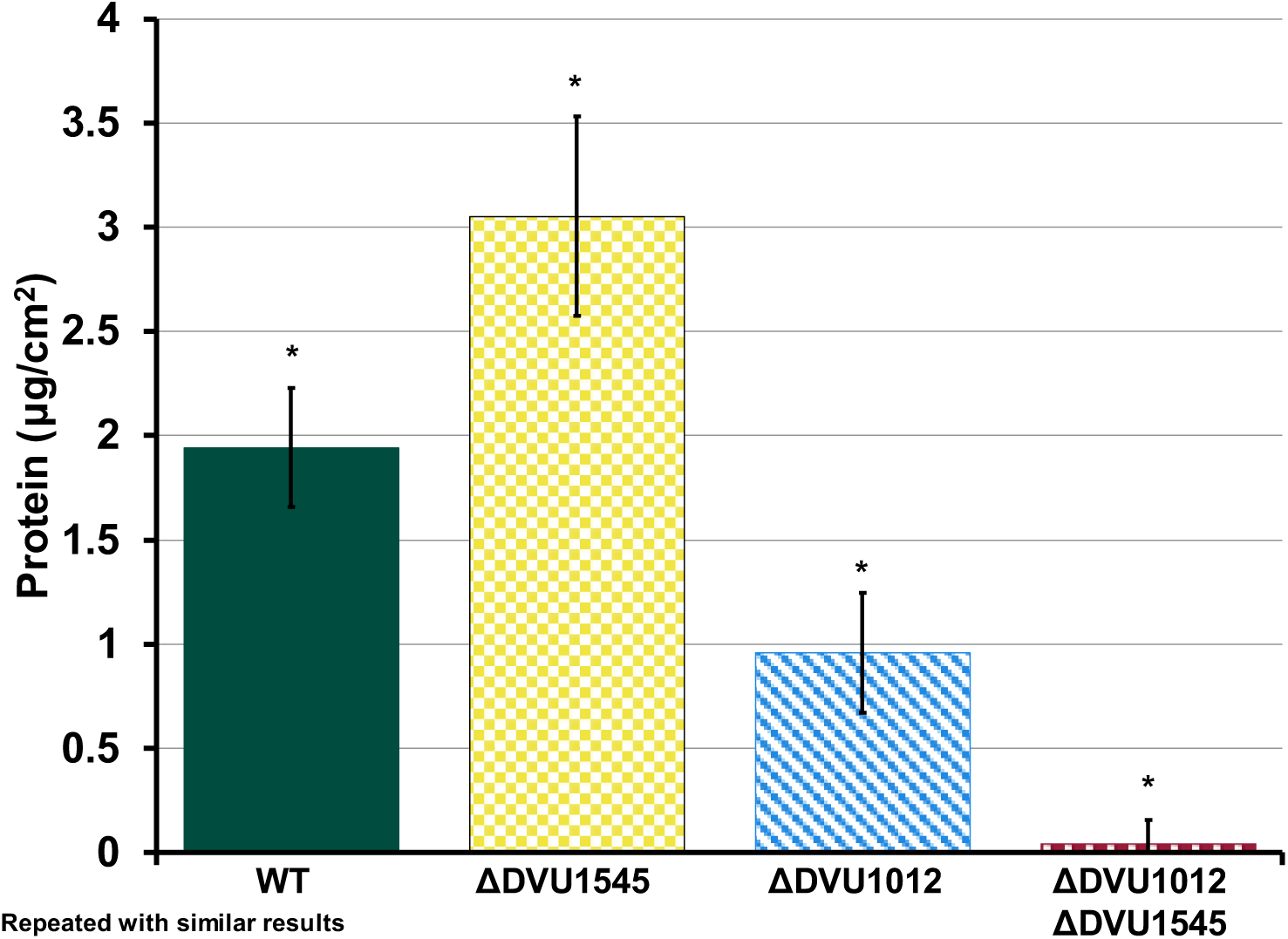
Early biofilm growth under static conditions for wild type and the biofilm adhesin mutants. Protein (µg/cm^2^) of biofilms formed by WT and the biofilm adhesin deletion mutants. Error bars indicate standard deviation across replicates (n=3). Asterisks above data points denote statistical significance as determined by a one-way ANOVA and Tukey Kramer post-hoc test (p < 0.01).

*D. vulgaris* strain DP4 (formerly strain DePue) is highly similar to *D. vulgaris* Hildenborough, having an average nucleotide identity of 98.77% (16, 17), but a homolog for DVU1545 is missing in this strain. The same experiment for biofilm formation under static culture conditions was performed with WT, ΔDVU1545, and strain DP4 to determine whether the closely related strain formed comparable biofilm biomass to the ΔDVU1545 strain. DP4 formed more biofilm than *D. vulgaris* Hildenborough WT but less biofilm than the ΔDVU1545 strain and was not significantly different from either (Figure 4). Nevertheless, this supported the observation that the absence of DVU1545 may result in more biofilm biomass in early biofilms under these conditions.

**Figure 4.**
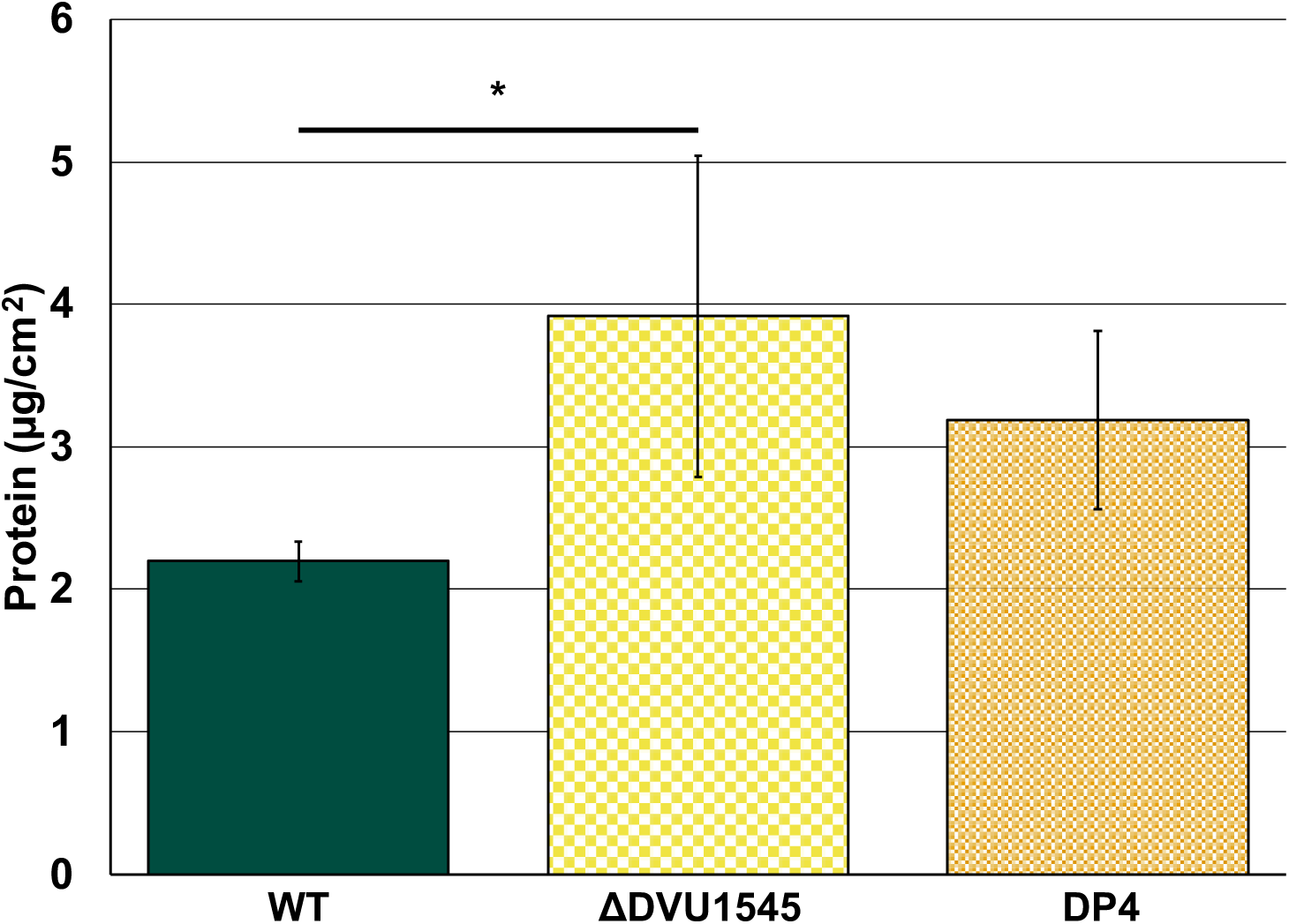
Early biofilm growth under static conditions for *D. vulgaris* strains Hildenborough (WT and ΔDVU1545) and DP4. Error bars indicate standard deviation across replicates (n=3). Line and asterisk above samples indicate statistically significant differences as calculated with a one-way ANOVA and Tukey-Kramer post hoc test (p < 0.05).

### Absence of biofilm proteins DVU1012, DVU1545 affect surface colonization and cell aggregation

Real-time microscopy was used to image biofilm formation over time within recirculating capillary reactors for WT and the deletion mutants to determine differences in surface colonization between the strains. Cultures were recirculated at a rate corresponding to the relative shear force experienced when cultured within the CDC bioreactors (18). Images were taken every two minutes for 48 h, with five positions imaged in each capillary reactor across three separate capillary reactor experiments per strain. Monitoring biofilm formation in real time revealed differences in cell strategy and surface colonization between WT and the biofilm protein deletion strains (Figure 5). WT and ΔDVU1545 cultures had higher proportions of cell aggregation compared to ΔDVU1012 and the double deletion mutant cultures (Figure 5a). Cells in WT cultures more often attached as cell aggregates rather than single cells. In contrast, the ΔDVU1012 cultures showed little aggregation, and many observed attachment events were by single cells (Figure 5a). While monitoring attached ΔDVU1012 cells over time, few microcolonies formed, though the cells appeared to be alive and were likely capable of replication.

**Figure 5.**
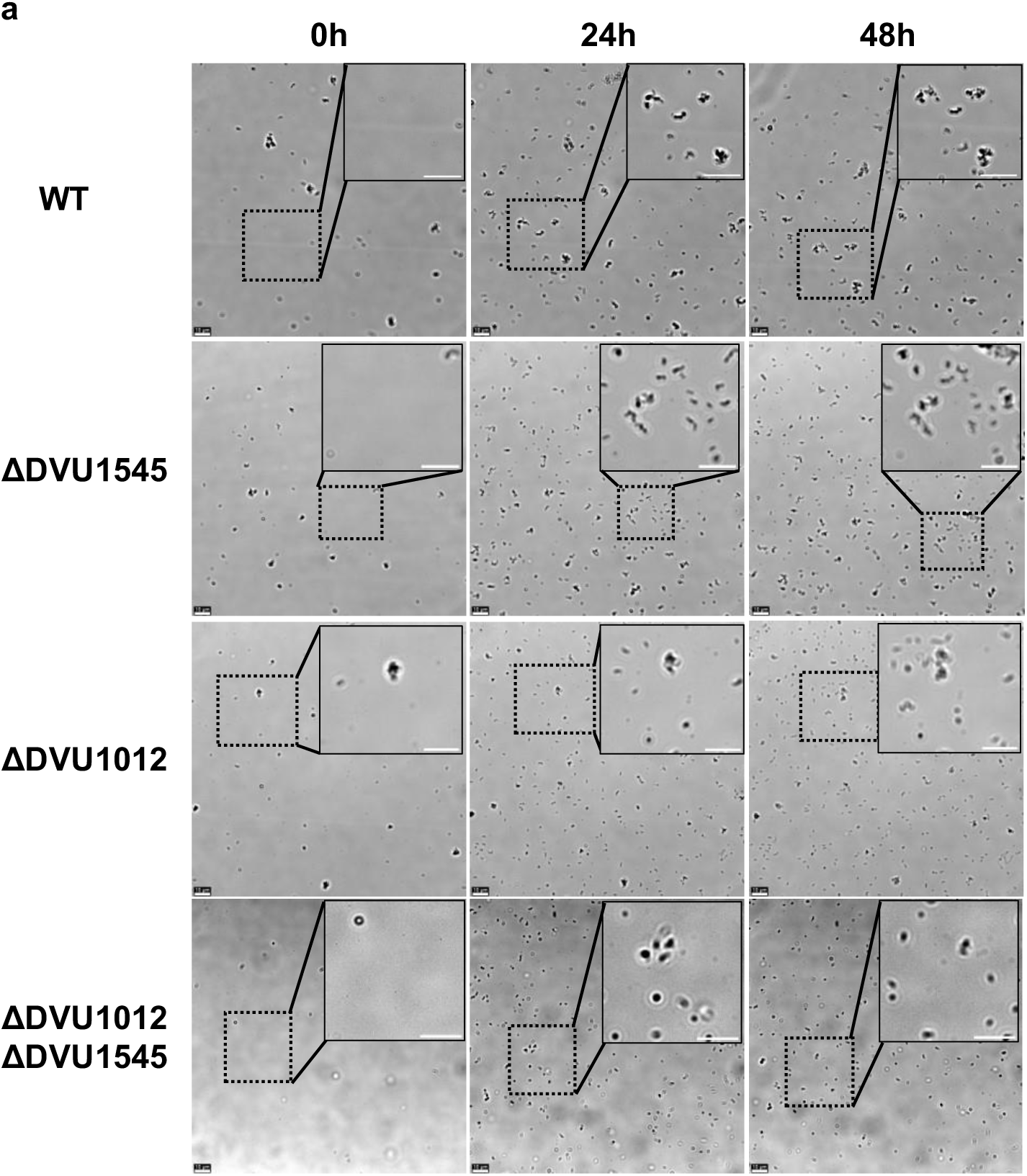
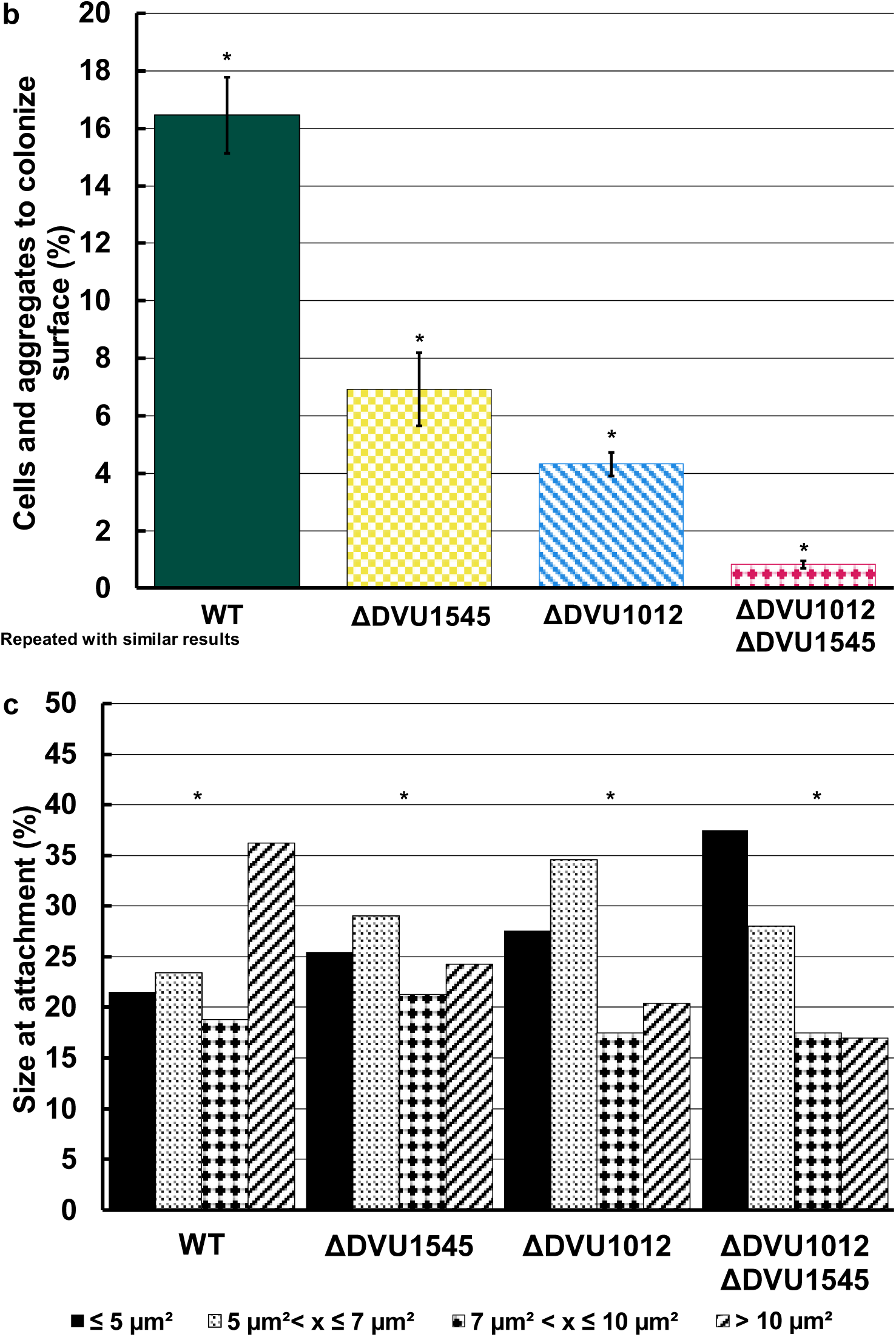
Differences in surface attachment and cell strategy of wild type and the biofilm adhesin mutants. Images of cells on the glass surface are shown for WT and the biofilm adhesin mutants at 0, 24, and 48 hours post-inoculation (a). The scale bar in bottom right of images represents 10 µm. The percentage of cells and cell aggregates to become surface-colonized (b). Error bars indicate standard deviation across replicate positions (n=5). Asterisks indicate statistical significance from a one-way ANOVA and Tukey Kramer post-hoc test (p < 0.05). The average number of cells and cell aggregates tracked during this experiment +/- standard deviation for these data are as follows: WT, 590 +/- 29; ΔDVU1545, 1958 +/- 181; ΔDVU1012, 2034 +/- 442; and ΔDVU1012 ΔDVU1545, 3630 +/- 933. Size distribution of surface-attached cells and cell aggregates at initial attachment for WT and biofilm adhesin mutants (c). The four size ranges selected are the quartiles for the entire dataset. Asterisks indicate statistical significance calculated using the Kruskal Wallis test and Conover post-hoc test (p < 0.05).

By tracking cells and cell aggregates throughout the experiment, the time that they remained attached was measured to differentiate reversibly attached cells from irreversibly attached cells (those considered to be surface colonized). We defined surface colonization as cells and cell aggregates that came into frame after the start of the experiment and remained in one position for at least 16 h. This length of time was determined by comparing the percentages of surface colonization at different lengths of time in increments of 1 h. We found that after 16 h the change in surface colonization percentages minimally changed, meaning if the cell or cell aggregate attached for 16 h, it was likely to remain attached for the duration of the experiment. So, we set the minimum time for defining colonization at this time. A significantly less proportion of tracked cells and cell aggregates of single deletion mutants colonized the surface compared to WT under these conditions (Figure 5b). The absence of DVU1012 caused an even greater reduction in surface colonization than the absence of DVU1545. The double deletion mutant strain lacking both DVU1012 and DVU1545 showed minimal surface colonization. The size of cells or cell aggregates that colonized the glass surface was measured at the time of attachment by particle tracking software (Figure 5c). WT cultures had the highest proportion of cell aggregates greater than 10 µm^2^ when first attached. The ΔDVU1012 cultures had lower proportions of attached aggregates greater than 7 µm^2^, and the double deletion mutant had few attachment events and, of those that attached, most were <5 µm^2^ and likely were single cells. Thus, surface colonization was impacted by the loss of either DVU1012 or DVU1545 and DVU1012 was important for cell aggregation and microcolony formation.

## Discussion

Biofilm adhesins provide a broad range of functions in the biofilm life cycle for bacteria and eukaryotic species, from surface colonization to biofilm maturation (19). Some species have multiple biofilm adhesins, and these proteins fulfill different roles and show differences in expression (11, 12). For example, biofilm adhesins, LapA and LapF, in environmental pseudomonads are important for surface colonization and cell-cell aggregation, respectively (11, 12). Mutants of *lapA* in the species *P. fluorescens* had a significant decrease in overall biofilm formation and surface colonization, whereas a mutation of *lapF* did not result in a complete loss of the phenotype but did lead to a decrease in overall biofilm formation and reduction in cell-cell attachment. We previously identified two proteins in the sulfate-reducing bacteria *D. vulgaris* Hildenborough, DVU1012 and DVU1545, localized to the cell surface by a type I secretion system and with similar sequence motifs to LapA and LapF, respectively (10). However, our single deletion mutants of DVU1012 or DVU1545 did not have a loss in the biofilm phenotype and had similar biofilm formation rates to WT (10). The double deletion mutant was deficient in biofilm formation; thus, at least one of these biofilm proteins is required for biofilm protein, but either adhesin is sufficient. The purpose of this current study was to determine differences in biofilm protein composition between the WT and single deletion strains, as well as determine the differences in cell strategy and surface colonization by WT, the single deletion mutants, and the double deletion mutant.

Proteomic analyses showed that the relative abundance of proteins DVU1012 and DVU1545 were not significantly different between WT and the single deletion mutants (Figure 2a). The protein DVU1012 was approximately four times more abundant than DVU1545 which is similar to what has been reported previously (9, 20).The protein DVU1012 was previously found as the most abundant protein within the extracellular matrix of biofilms formed by *D. vulgaris* (9) and DVU1012 was ranked 80^th^ in our whole-cell proteomics data and the most abundant protein predicted to be extracellular. This indicates that the absence of one biofilm protein does not lead to an increase in the other protein’s abundance, and that the proteins do not directly compensate for the other within the biofilm matrix. Yet, there is no reduction in whole biofilm biomass, measured as protein, between the strains when the gene is removed from the bacterium. Based on the proportion of the protein DVU1012 within the biofilm proteome of WT, we estimated that the protein DVU1012 abundance to be 0.14 µg/cm^2^. When comparing biofilm mass coverages of biofilms between WT and ΔDVU1012, this falls within the standard deviation of both strains. From the proteomics data, we did not determine a clear change in biofilm matrix protein composition between the strains except for the absence of the deleted adhesin in each single deletion mutant and a hypothetical protein encoded by DVU2135 that was increased in the ΔDVU1012 strains and was predicted to localize to the extracellular matrix, but its function is unknown. Many of the proteins that did significantly change in abundance were associated with cell wall structure and chemotaxis. The proteins with the largest increase in abundance within biofilms of ΔDVU1012 strain compared to WT are predicted to be involved in cell wall structure and in chemotaxis (encoded by DVU2365 and DVU2075, respectively). (Figure 2b). The protein encoded by DVU2365 has 51 % identity to the UDP-2,3-diacylglucosamine diphosphatase, LpxI, from *D. ferrophilus* IS5, which is involved in lipid biosynthesis and is increased in expression within biofilms of *D. ferrophilus* IS5 on carbon steel surfaces (21). Interestingly, the proteins with the largest decrease in abundance within biofilms of the ΔDVU1012 strain are also predicted to be involved in the cell wall structure. For example, the proteins encoded by DVU1202, DVU3046, and DVU0336 are predicted to play a role in cell wall biosynthesis. It is possible that the absence of the protein DVU1012, which localizes to the cell surface and has been previously shown to be the most abundant protein within the biofilm matrix (9), leads to a change in the cell surface and cell wall compositions.

A similar trend in changed abundances occurred biofilms of the ΔDVU1545 strain compared to WT. The proteins encoded by DVU0788 and DVU1884 are a rod-shape determining protein MreC and methyl-accepting chemotaxis protein, respectively, showed the highest increase in abundance in biofilms of the ΔDVU1545 strain and are also predicted to be involved in cell shape and chemotaxis. The chemotaxis protein encoded by DVU1884 is predicted to contain a PAS domain, which has been linked to biofilm regulation in other species such as *Pseudomonas aeruginosa* (22). The protein with the largest decrease in abundance, encoded by DVU3387, has predicted domains related to glycosyl hydrolase enzymes and thus also related to cell wall synthesis. These changes in abundance of proteins related to cell wall, cell partitioning, and chemotaxis for both single deletion mutants suggests that there are additional changes in cell surface composition in the absence of either biofilm adhesin. Both adhesion proteins are abundant within WT cells, and so their absence likely requires a change in the cell surface composition to compensate and maintain the integrity of the membrane.

While measuring biofilm formation over time under different growth conditions, the rates of biofilm maturation in the WT and single deletion mutants was similar in the CDC reactors (Figure 1), as has been shown previously (10). However, the measurement of early biofilms cultivated under static conditions did reveal biofilm phenotypic differences between the strains. The absence of protein DVU1012 caused a significant decrease in biofilm under static conditions, whereas the inverse was seen in the absence of protein DVU1545 (Figure 3). The double deletion strain had little to no measurable biofilm biomass. Thus, the presence of at least one biofilm adhesin is also required for biofilm formation under static conditions cultures. *D. vulgaris* strain DP4 is highly similar to *D. vulgaris* Hildenborough genetically. Both genomes are highly syntenous with most of the variability being the presence of different phage islands and different CRISPR spacers; but strain DP4 is missing a homolog for DVU1545 (16). We compared early biofilm formation of DP4 to our strains of *D. vulgaris* Hildenborough to determine whether a native strain would also have increased biofilm when DVU1545 was missing. DP4 formed more biofilm when compared to *D. vulgaris* Hildenborough WT and less than that of the ΔDVU1545 strain, though it was not significantly different from either (Figure 4). This increase in biofilm by the ΔDVU1545 strain was not observed under shear conditions in the CDC reactors (Figure 1) or capillary reactors (Figure 5). We observed more cell attachment events in capillary reactors with the ΔDVU1545 strain than the WT strain, but more of these cells were reversibly attached and did not colonize the surface. Thus, the absence of DVU1545 may impact resistance to shear and/or a transition from reversible to irreversible attachment.

Real-time monitoring of biofilm formation by the strains revealed differences in cell strategy and surface colonization by WT and biofilm protein deletion mutants. Each capillary was inoculated with the same cell density of 1 x 10^6^ cells/mL, so differences seen in the number of surface colonized cells and cell aggregates would be due to the absence of one or both proteins. We found the absence of either biofilm adhesin caused a decrease in surface colonization compared to WT, with a greater effect seen in the absence of DVU1012 (Figure 5b). This indicates the loss of protein DVU1012 affects surface colonization more so than the protein DVU1545. Results from comparing the proportion of sizes of tracked objects (cells and cell aggregates) when first attached showed WT cultures had the highest proportion of cell aggregates compared to the other strains and most often attached as cell aggregates (Figure 5c). Cells lacking DVU1545 had a more even distribution of particle sizes and attached as both individual cells and cell aggregates. Cells lacking DVU1012 tended to attach more often as single cells, and few colonized cells formed microcolonies over time. The attached cells were likely active and capable of cell division, but the daughter cells did not remain on the surface or did not remain adjacent to the parent cell upon division. This suggests that biofilms of cells lacking DVU1012 have more confluent growth across the surface that eventually fills the surface area to then establish a mature biofilm.

Overall, we conclude that the absence of either biofilm adhesin does not cause a proportional change in the abundance of the other biofilm adhesin, nor does it cause proportional changes in other proteins that show potential to compensate within the biofilm matrix. The absence of DVU1012 impacts cell aggregation and the absence of DVU1545 leads to phenotypic differences depending on shear conditions. This work has further illuminated the knowledge of biofilm adhesins in *Desulfovibrio* species and showed differences in biofilm adhesin functionality compared to as the Lap system in *Pseudomonas* spp. We have shown physiological differences between the two proteins, DVU1012 and DVU1545, and how they differ from the proteins LapA and LapF, important for surface attachment and cell attachment, respectively, in environmental pseudomonads. Recently, Karbelkar et al. 2024 compared the predicted biofilm regulatory components to those of *P. fluorescens* (13). Like the soluble, periplasmic protease, LapG, the protease encoded by DVU1019 in *D. vulgaris* Hildenborough cleaves DVU1012 at a di-alanine motif (106AA107). However, a transmembrane (TM) helix in the protein DVU1019 is proposed to anchor DVU1019 to the inner membrane and is essential for function. LapD binds cyclic-di-GMP by a catalytically inactive c-di-GMP phosphodiesterase domain (EAL) (23). Under high c-di-GMP concentrations LapD binds LapG and sequesters it from cleaving LapA. Under low c-di-GMP concentrations, LapG is released and cleaves LapA, releasing it from the cell surface and allowing the cell to detach from the surface (24). Karbelkar et al. 2024 showed that a similar protein, encoded by DVU1020, contains a catalytically inactive phosphodiesterase domain, HD-GYP, instead of EAL domain, but has a high affinity for c-di-GMP and modulates DVU1019 protease activity, similar to the LapD-LapG complex. How this complexation is impacted by DVU1019 being bound to the inner membrane remains unknown. For simplicity, Karbelkar et al. 2024 referred to these proteins as DvhA-F analogous to LapA-F. Our current study and these studies have revealed differences in these biofilm systems. Given these differences, we propose that these proteins be named for their function in the biofilm formation system (Bfs). We propose the following naming: BfsA for the adhesin encoded by DVU1012; BfsEBC for DVU1013, DVU1017, and DVU1018 encoding the type I secretion system components TolC, ATP transporter, and membrane fusion protein; BfsG for the protease encoded by DVU1019, BfsD for the effector protein encoded by DVU1020; and BfsF for the adhesin DVU1545. These subunits maintain the connection between the Bfs and Lap systems but acknowledge the functional differences between these systems and will facilitate further comparative studies.

## Materials and Methods

### Strains, media, and growth conditions

The construction and storage of the parental strain (Δ*upp*; JWT700; used as wild type in this study) and the biofilm adhesin mutants ΔDVU1012 (JWT705), ΔDVU1545 (JWT706), and double-deletion mutant (ΔDVU1012 ΔDVU1545; JWT709) used in this study have been described previously (10). The bacterium *D. vulgaris* DP4 was provided by Judy Wall and grown under the same conditions as the *D. vulgaris* Hildenborough strains. Cultures were prepared in MOYLS4; LS4D with 60 mM sodium lactate and 50 mM sodium sulfate (60:50); or LS4D with 20 mM sodium lactate, 16.6 mM sodium sulfate, and 106 mM sodium chloride (20:16.6), as described previously and indicated where relevant below (10). All incubations were done at 30 °C unless otherwise indicated. Starter cultures were prepared by inoculating 1 ml of a frozen stock into 10 ml of anoxic MOYLS4 within a sealed Balch-type 18 x 150 mm tube with a nitrogen headspace. These were incubated until the cultures reached an optical density at 600nm (OD600) of approximately 0.8 (mid- to late-log phase) measured with a Genesys 30 Visual Spectrophotometer (Thermo Scientific, Waltham, MA, USA). Tubes for culturing biofilms under batch and static conditions were prepared by placing a halved borosilicate glass slide (Fisher Scientific, Hampton, NH, USA) sitting obliquely in 10 ml of 60:50 LS4D within an anoxic Balch tube and sterilized by autoclaving. Cultures were inoculated at a 1:50 dilution of the starter culture and incubated for 24 h. At the end of each experiment, to check for aerobic contamination of cultures, 10 µL of each culture was inoculated onto LC+glucose agar plates, prepared as described previously (10), and incubated at the same temperature as the experimental conditions (either 30 °C or 23 °C).

### Biofilm formation under continuous culture and shear conditions

To determine differences in biofilm protein composition, strains WT, ΔDVU1012, and ΔDVU1545 were cultivated within the CDC bioreactors (Biosurface Technologies, Bozeman, MT, USA) as done previously (10). Briefly, starter cultures of WT, ΔDVU1012, and ΔDVU1545 were used to inoculate 20:16.6 LS4D media within the CDC bioreactors and incubated at 30 °C. Reactor headspaces were continuously sparged at 100 ml/min with nitrogen gas passed through a sterile gas filter. Once the cultures within the bioreactors reached an OD_600_ of ∼0.7, media was flowed into each reactor at 0.7 ml/min. Biofilm and planktonic samples were taken daily for five consecutive days. At 72 hours, biofilm samples were taken in triplicate for proteomic analyses. All biofilm and planktonic samples were pelleted via centrifugation (17000 x g, 2 minutes) and stored at -20 °C. Planktonic samples from 120 hours were used for genomic DNA extraction using the Wizard Genomic DNA Purification Kit, (Promega, Madison, WI, USA). Genotypic confirmation of each strain was done using PCR as previously described (10). Protein from biofilm pellets was quantified via the Bradford Assay (25).

### Biofilm formation under batch culture and static conditions

Strains of DvH were grown from freezer stock until reaching late log phase of growth (OD_600_ ∼0.8). Strains were then sub-cultured in a 1:50 dilution into anoxic Balch tubes containing 60:50 LS4D medium either with or without a halved glass slide in triplicate and incubated at 30 °C under static conditions. OD_600_ measurements were taken over time to track planktonic growth over time. After 24 hours of growth, biofilm samples were taken from each strain as reported previously (10). Biofilm samples were then stored at -20 °C until quantified via the micro assay protocol of the Bradford Assay (25, 26).

### Capillary Flow Cell Imaging

Square glass capillary tubes (Friedrich & Dimmock, Millville, NJ, USA) measuring 1 x 1 x 100mm with an average wall thickness of 0.17mm were cleaned with 95% EtOH and connected to polypropylene-derived tubing (Avantor, Radnor Township, PA, USA). Capillary and Masterflex^®^ L/S^®^ 13 Norprene^TM^ tubing (Avantor Inc., Radnor, PA, USA) were sterilized via autoclave (121 °C, 30 minutes), and then aseptically connected. The capillaries were then placed within a 3D-printed holder and then secured onto the mechanical stage of a DMi8 epifluorescence microscope (Leica, Deerfield, IL, USA).

Once cultures showed signs of active growth in 60:50 LS4D, samples of WT, ΔDVU1012, ΔDVU1545, and ΔDVU1012 ΔDVU1545 were used for direct cell counts in a Neubauer counting chamber (Levy Ultra Plane; Clay-Adams Co., New York, USA) and the cultures were diluted to 1*10^6^ cells/ml with 60:50 LS4D. Thirty milliliters of each culture were then aseptically transferred to the capillary flow cell system and then recirculated through the tubing and capillaries at 300 µl/min for 10 minutes to homogenize the cultures within the tubing. A Masterflex^®^ L/S^®^ Digital Drive with a standard pump head for L/S 13 tubing was used to recirculate the cultures at 0.002 ml/min within the flow cell system. During this time, five positions were chosen at random on each capillary tube for imaging. After 10 minutes, flow was reduced to 2 µl/min for image acquisition. Microscopy images were taken through a HC PL FL L 63x/0.70 CORR PH2 objective lens (NCI, Brooklyn Park, MN, USA) and a K5 microscope camera (Thomas Scientific, Chadds Ford Township, PA, USA). Phase contrast images of each inner capillary surface were taken every 10 minutes for 48 hours at each position, with autofocus correction occurring every 1 hour. Image series were then saved for processing and analysis. Image processing and analysis was performed using Leica Application Suite X v.3.7.4.23463 (LAS X) to determine attachment events for cells and cell aggregates, and cell and cell aggregate sizes. Object size and location data were used to track particles across each image series with LAS X.

### Peptide sample preparation

Proteins were extracted from biofilm samples as done previously with modifications noted here (27, 28). Briefly, biofilm pellets were resuspended in 1ml lysis buffer (10mM Tris-HCl pH 8.0, 1% w/v SDS, 0.1M dithiothreitol (DTT)) and incubated at 95 °C for 30 minutes. Biofilm samples were then sonicated on ice (20% amplitude, 30 second intervals, five intervals) using a Branson 450 digital sonifier with a 102C converter (Branson Electronic Corp, St. Louis, MO, USA). Lysed biofilm samples were then centrifuged at 14000 x g and 4 °C for 10 minutes. Supernatants were transferred to new tubes before adding trichloroacetic acid to a final concentration of 25% (v/v) and stored at 4 °C overnight to precipitate solubilized proteins. Samples were then pelleted via centrifugation (20800 x g, 10 min), the supernatant discarded, and the pellets washed three times with chilled (∼4 °C) acetone. Protein pellets were solubilized in guanidine buffer (6M guanidine HCl, 10mM DTT in Tris CaCl_2_ buffer (50mM Tris, 10mM CaCl_2_, pH adjusted to 7.6 with HCl)) at 60 °C, 1500 rpm for one hour before mixing with UA buffer (8 M urea in 0.1M Tris-HCl pH 8.5) in a 1:3 ratio and concentrated on a 30 kDA filter (MilliporeSigma, Burlington, MA, USA) via centrifugation (14000 x g, 15 min). Columns were washed with UA buffer, IAA buffer (50 mM iodoacetamide in UA buffer), and then ABC buffer (50 mM NH_4_HCO_3_) via centrifugation as done in the previous step. Concentrated protein extracts were then digested with 3 µl trypsin according to manufacturer’s instruction (Promega, Madison, WI, USA) at 37 °C overnight. Digested protein samples were then eluted with ABC buffer and desalted using the Pierce C-18 Spin Columns according to the manufacturer instructions (Thermo Scientific, Waltham, MA, USA). Peptide samples were then immediately quantified using the Pierce Quantitative Colorimetric Peptide Assay (ThermoScientific, Waltham, MA, USA). Peptides were then frozen at -80 °C, then lyophilized at -50 °C and 0.015 mBar for six hours. Lyophilized peptides were then sent to the IDeA National Resource for Quantitative Proteomics (Little Rock, AR, USA) for proteomic analyses.

### Proteomic Analysis

Tryptic peptides were separated from other peptides with reverse phase XSelect CSH C18 2.5 µm resin (Waters, USA) on an in-line 150 x 0.075 mm column using an UltiMate 3000 RSLCnano system (ThermoScientific, Waltham, MA, USA). Separated peptides were then eluted using a 90-minute gradient from 98:2 to 65:35 buffer A:B ratio (Buffer A: 0.1% formic acid, 0.5% acetonitrile; Buffer B: 0.1% formic acid, 99.9% acetonitrile). Eluted peptides were ionized by electrospray at 2.4 kV followed by mass spectrometric analysis on an Orbitrap Eclipse Tribrid mass spectrometer (ThermoScientific, Waltham, MA, USA). Mass spectrometry (MS) data were acquired using the FTMS analyzer in profile mode at a resolution of 120,000 over a range of 375 to 1200 m/z. Following HCD activation, MS/MS data were acquired using the ion trap analyzer in centroid mode and normal mass range with a normalized collision energy of 30%. Proteins were identified using Sipros v1.1 (29) and quantified using ProRata v1.1 (30). Briefly, Sipros Ensemble was used to search mass spectrometry spectra against the proteins of *D. vulgaris* Hildenborough acquired from https://microbesonline.org. Raw search results were filtered to achieve a 1 % false discovery rate at the peptide level, which was estimated using the target-decoy approach. Peptide identifications were assigned to protein or protein groups in accordance with the parsimonious rule. Protein quantification was achieved through intensity-based label-free analysis using ProRata. Protein abundances were quantified by the total peak height of all quantified peptides from a protein, normalized against the average total of all data sets. Relative fold change analysis of identified protein intensities was done using Excel and methods derived from Aguilan et al (31). Briefly, protein intensities were normalized before determining fold change differences between WT and biofilm adhesin mutants. An F-test was performed to determine whether the normalized protein intensities were homoscedastic or heteroscedastic, and then a Student’s t-test with independent samples was performed either using the homoscedastic or heteroscedastic selection. The cutoff for significant fold changes was set to either positive or negative two, and p < 0.05. Localization of proteins from *D. vulgaris* Hildenborough was predicted using PSORTb v.3.0.3 (32). Functional categories of proteins were determined using the Cluster of Orthologous Groups (COGs) database (33).

## Data Availability

The raw mass spectrometry data from the proteomics analysis have been deposited in the EMBL-EBI Proteomics Identifications Database (PRIDE) under the doi: (doi will be made available upon acceptance or request).

## Acknowledgements

The culture of *D. vulgaris* DP4 was generously provided by Dr. Judy D. Wall, University of Missouri. Special thanks to the proteomics core at the University of Arkansas Medical Center and the IDeA National Proteomics workshop, for the training on the workflow and analyses for proteomics. Thanks to Dr. Peter J. Walian at Lawrence Berkeley National Lab for his expertise and assistance with the methods for proteomics of *Desulfovibrio* species. The funding for this work was provided by the University of Oklahoma School of Biological Sciences.

